# A novel co-segregating *DCTN1* splice site variant in a family with Bipolar Disorder may hold the key to understanding the etiology

**DOI:** 10.1101/354100

**Authors:** André Hallen, Arthur J.L. Cooper

**Affiliations:** Department of Molecular Sciences, Macquarie University, Sydney, Australia, NSW 2109; Department of Biochemistry and Molecular Biology, New York Medical College, Valhalla, NY 10595

**Keywords:** Bipolar disorder genetics, DCTN1, p150Glued, retrograde axonal transport, autophagy

## Abstract

A novel co-segregating splice site variant in the *Dynactin-1* (*DCTN1*) gene was discovered by Next Generation Sequencing (NGS) in a family with a history of bipolar disorder (BD) and major depressive diagnosis (MDD). Psychiatric illness in this family follows an autosomal dominant pattern. *DCTN1* codes for the largest dynactin subunit, namely p150^Glued^, which plays an essential role in retrograde axonal transport and in neuronal autophagy. A GT→TT transversion in the *DCTN1* gene, uncovered in the present work, is predicted to disrupt the invariant canonical splice donor site IVS22+1G>T and result in intron retention and a premature termination codon (PTC). Thus, this splice site variant is predicted to trigger RNA nonsense-mediated decay (NMD) and/or result in a C-terminal truncated p150^Glued^ protein (ct-p150^Glued^), thereby negatively impacting retrograde axonal transport and neuronal autophagy. BD prophylactic medications, and most antipsychotics and antidepressants, are known to enhance neuronal autophagy. This variant is analogous to the dominant-negative *GLUED Gl*^*1*^ mutation in *Drosophila* which is responsible for a neurodegenerative phenotype. The newly identified variant may reflect an autosomal dominant cause of psychiatric pathology in this affected family. Factors that affect alternative splicing of the *DCTN1* gene, leading to NMD and/or ct-p150^Glued^, may be of fundamental importance in contributing to our understanding of the etiology of BD as well as MDD.

## 1. Introduction

Bipolar disorder (BD) is a chronic and severe psychiatric disorder. In most cases, it is a recurrent psychiatric disorder characterized by oscillations between mania and major depressive episodes, although some patients may only experience mania. Mania in affected patients may or may not involve psychosis. BD is subdivided into two main subtypes, notably type I (BDI) and type II (BDII). BDI is generally regarded as more severe than is BDII due to the presence of mania in BDI patients compared to an absence or decreased severity of mania in BDII affected individuals [1]. Cyclothymic disorder or bipolar disorder III (BDIII) is regarded as part of the BD spectrum of illnesses, sharing similar characteristics with other BDs. However, BDIII is rarely diagnosed because symptoms are generally below the threshold required for a definitive diagnosis of BD. Nevertheless, in many cases individuals with cyclothymic disorder also go on later to experience episodes of mania and develop BDI [2]. BD often requires repeated hospitalizations and is an expensive illness to treat both for the affected individual and for society, often also resulting in significant unemployment [3]. The disorder is often associated with considerable morbidity and a substantial increased risk of suicide (10-20%) [4]. BD is considered a complex illness, often suspected to involve interactions between genetic and environmental factors such as stressful circumstances as well as seasonal changes [5,6].

Our present understanding of the underlying molecular basis of BD has thus far been insufficient to explain its etiology and there are no existing biochemical- or genetic-based laboratory tests available to validate a diagnosis. Currently, the diagnoses are based on observations and history taking, and thus are not predictive of illness risk for individuals who may later develop illness. BDI and BDII have a lifetime world-wide prevalence of ∼0.6%, and ∼0.4% respectively, and the lifetime prevalence of sub-threshold BD has been estimated to be ∼1.4%% [7]. Major depressive diagnosis (MDD), on the other hand, is more common with a lifetime world-wide prevalence of ∼4-5% [8]. BDI has long been known to be highly heritable (> 80 %) [9]. As major depressive episodes are a notable feature of BD, it is not surprising that MDD is also a common feature in extended families with BD, implying a common etiology in these affected families[10]. The age of onset of mania in BDI patients shows a peak in early adulthood in a 35 year UK study [11]. These findings are corroborated by a large study in the Netherlands where the authors found two peaks in the age of onset for BD, one likewise in early adulthood (mainly for BDI) and another peak later in life [12]. Similarly, MDD patients having a family history of MDD, and experiencing recurrent depressive episodes, also exhibit a peak for age of onset in early adulthood [13].

Many genes and chromosomal regions have been implicated in BD. However, to this date no single gene defect alone has been determined to be causative [14,15]. One of the primary difficulties in discovering candidate disease-causing variants is in defining a homogeneous cohort of patients. In this research study we have attempted to address this problem by focusing on a single core family with two siblings diagnosed with BDI, a parent with BDIII and a family pedigree which strongly suggests that psychiatric illness follows an autosomal dominant inheritance pattern (Fig. 1). Both BDI siblings have a decades-long psychiatric history and have experienced multiple manic episodes, with psychotic features, and depressive episodes satisfying the DSM-5 (Diagnostic and Statistical Manual of Mental Disorders – Fifth Edition) criteria for BDI. It is known that there is a high risk of psychiatric disorders among the relatives of patients with BDI and MDD [16,17]. Thus, we did not discriminate among different psychiatric diagnoses in the extended family as the family pedigree historically includes individuals diagnosed with BDI, MDD, or BDIII (Fig. 1). Apart from the three affected subjects, there were no other living individuals with a psychiatric diagnosis in the extended family. Genomic data from twelve related controls, and one unrelated control, were also used to filter variants to assist in the discovery of the novel co-segregating *DCTN1* splice site variant described in this research.

**Figure 1.**
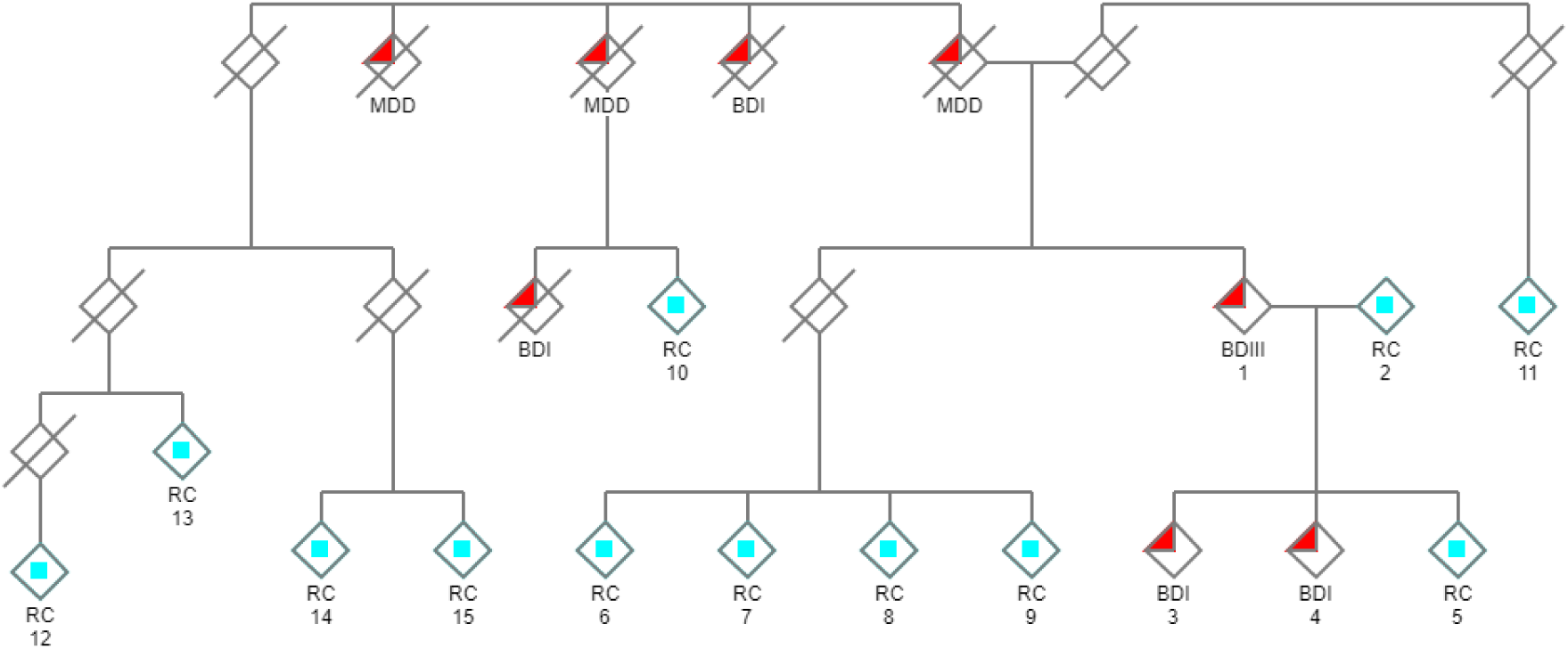
Pedigree of the family under investigation. Individuals with a psychiatric history are denoted with filled red triangles. Psychiatric illness in this extended family is inherited in a manner resembling autosomal dominance. WES was performed and subtractive analysis was used to discover novel co-segregating candidate gene variants in the three living affected individuals in the core group (#1,3,4), using related unaffected living relatives as negative controls (related controls/RC: light blue squares). (/ = deceased; and males and females are not distinguished for reasons of privacy).

## 2. Materials and Methods

### 2.1 Genome sequencing

Whole exome sequencing (WES) was used in the present study, which included genetic analyses of three affected subjects, twelve related unaffected controls (Fig. 1), and one unrelated unaffected control. This research project was approved by the Macquarie University Human Ethics Committee: “Bipolar Disorder I: a family-based genome sequencing study” (Ref#: 5201400393). In accordance with standard ethical practice all participants who were genetically tested provided their signed informed consent. Genomic DNA was extracted from whole blood samples using accepted published protocols and submitted to the relevant sequencing center for sequencing (Macrogen, South Korea). 100 bp Paired-end reads were used. Agilent SureSelect All Exon V5 kit was used for NGS exome enrichment, and NGS was performed using Illumina HiSeq 4000 (Illumina, San Diego, USA) at an average depth of 100x. Illumina sequencing files were deposited in the NCBI SRA repository (accession: PRJNA607165).

### 2.2 Genomic alignment

All bioinformatic analyses were performed using the DNASTAR Lasergene Full Suite (DNASTAR, Madison, Wisconsin, USA, v17). Raw genomic data was aligned, and variants called and annotated using SeqMan NGen. Genomic raw FASTQ data was aligned to the GRCh37.p13-dbSNP150 genome template (pre-configured by DNASTAR) using the default low stringency layout option in order to maximize the true positive rate. A mer size of 21nt and minimum match percentage of 93% was used. The default alignment settings in the Seqman NGen application were used and duplicate reads were combined (minimum alignment length = 35, minimum layout length = 21, maximum gap size = 30). Reads were auto scanned for adaptor sequences and auto trimmed prior to alignment. The Agilent SureSelect All Exon V5 targeted region BED file was used covering 21,522 genes and 357,999 targeted exomes.

### 2.3 Analytical strategy

As the family pedigree suggests autosomal dominant inheritance is most likely (Fig. 1), the strategy used herein was to search for novel co-segregating variants because these variants would be the most likely to be disease-causing. This strategy has consistently been shown to have validity [18,19]. Effective use was made of numerous related controls to narrow the search for candidate genes by filtering out confounding variants also present in controls. Comparative analysis was performed using the ArrayStar application within DNASTAR Lasergene.

1. Variant calls were restricted to coding and splice regions, and a minimal variant call filter of the probability of not being reference ≥ 90% was applied.
2. Variant calls with a minor allele frequency (MAF) ≥ 0.01 in 1000 genomes phase 3 were filtered.
3. Functional prediction filtering was applied where only variants that were predicted to be deleterious in at least one of the following bioinformatic resources included in the ArrayStar application were retained; LRT [20], MutationTaster [20], or SIFT [21].
4. Using the Venn diagram feature of ArrayStar only co-segregating variants common to all three affected subjects and not present in any of the controls were retained.
5. The variant calls from the sequencing data of ∼120,000 exomes within the Genome Aggregation Database v2.02 (gnomAD) [22] were loaded into ArrayStar and all variant calls present in gnomAD were removed by filtering.

### 2.4 Bioinformatic predictions

In addition to bioinformatic predictive algorithms included in Arraystar, prediction of pathogenicity was also determined using the following bioinformatic resources: CADD [23], DANN [24], and FATHMM-XF [25]. Genomic evolutionary conservation was determined using GERP++RS [23], PhastCons100way_vertebrate [26] and PhyloP100way_vertebrate [26] from within ArrayStar. Predicted pathogenicity was also evaluated using Varsome which utilizes multiple bioinformatic algorithms [27].

### 2.5 Sanger sequencing validation

Samples #1-7 were further validated by Sanger sequencing. These samples covered all five members of the core family under investigation, including all three with a psychiatric diagnosis (Fig. 1, #1,3,4). As DNA samples were collected from different continents not all control DNA was available for Sanger sequencing. Quality/quantity of the extracted DNA was evaluated by gel electrophoresis and Qubit dsDNA BR Assay Kit (Life Technologies). To evaluate the probability of DNA degradation, gel electrophoresis was carried out by loading 10 µl (500 ng) of extracted DNA onto a 1% agarose gel. The following PCR primers were used: Forward-primer: 5’-TCATACTCCCCCTCCTGCAT-3’; Reverse-primer 5’-AATGAGGGGCTACTTGTGGC-3’. PCR was performed in a 100 µL volume, consisting of: 10 µL Qiagen PCR Buffer (10X), 2 µL of 10 mM dNTPs, primer sets, 0.5 µL of Taq DNA polymerase, and 4 µL (200 ng) of genomic DNA. The amplification program consisted of one initial denaturation at 94°C for 3 min followed by 30 cycles of 30s at 94°C for denaturation, 30s at 65°C for primer annealing, 30s at 72°C for extension and final extension at 72°C for 5 min. Products were separated on a 1% SYBR Safe agarose gel and visualized by Safe Imager™ 2.0 Blue Light Transilluminator (Thermo Fisher).

## 3. Results

Only a single novel co-segregating variant, predicted to be deleterious, was identified in the affected family (Fig. 1). When confounding variants were removed through filtering a transversion splice site variant in the *DCTN1* gene (IVS22+1G>T, GRCh37/hg19: 2:74,593,585C>A) was discovered using NGS (Table 1; suppl. material, S1), and validated by Sanger sequencing (Fig. 2, suppl. material, S2). It was the only novel variant, in coding or splice regions, that was found to co-segregate with the affected cohort and not be present in any of the controls (suppl. material, S1). This splice site variant is predicted to disrupt the canonical splice donor site at the invariant +1 position and result in retention of intron 22 and introduction of a subsequent PTC (Fig. A1 in appendix). The splice site variant is in a highly evolutionary conserved region (Table 2), and all functional prediction algorithms predicted the variant to be deleterious with no algorithms predicting the variant to be benign (Table 3, https://varsome.com/variant/hg19/2%3A74593585%3AC%3AA). There is a very rare alternative allele (n =1) noted at this coordinate in the dbSNP database (rs75942278) which contains a C>T transition. However, a transition (where nucleotide ring structure remains constant) is less likely to impair the spliceosome mechanism and result in donor splice site disruption compared to a transversion (where nucleotide ring structure changes).

**Table 1.** Variant counts for the *DCTN1* splice site variant [NC_000002.11: g.74593585C>A (GRCh37)] in affected subjects (#1,3,4); BDI, bipolar disorder I; BDIII, bipolar disorder III. Variant calls for all probands and controls are in suppl. material, S1.

| Subject # | Diagnosis | *DCTN1*<br>Count | *DCTN1*<br>Count |
| --- | --- | --- | --- |
|  |  | C | A |
| 1 | BDIII | 31 | 51 |
| 3 | BDI | 51 | 58 |
| 4 | BDI | 76 | 64 |

**Table 2.** Genomic evolutionary conservation data for the *DCTN1* splice site variant.

|  | GERP++RS | PhastCons100way Vertebrate | PhyloP100way Vertebrate |
| --- | --- | --- | --- |
| Score | 5.08 (range 1-6.18) | 1 (range 0-1) | 5.75 (range -20-10) |
| Conservation | Highly conserved | Highly conserved | Highly conserved |

**Table 3.** Functional predictions for the *DCTN1* splice site variant

|  | CADD | DANN | FATHMM-XF5 | MutationTaster |
| --- | --- | --- | --- | --- |
| Score | 25.6 | 0.9946 | 0.996 | 1.0 |
|  | [>20 = 1% most pathogenic] | [range 0-1] | [range 0-1] | [range 0-1] |
| Prediction | Deleterious | Deleterious | Deleterious | Deleterious |

**Figure 2.**
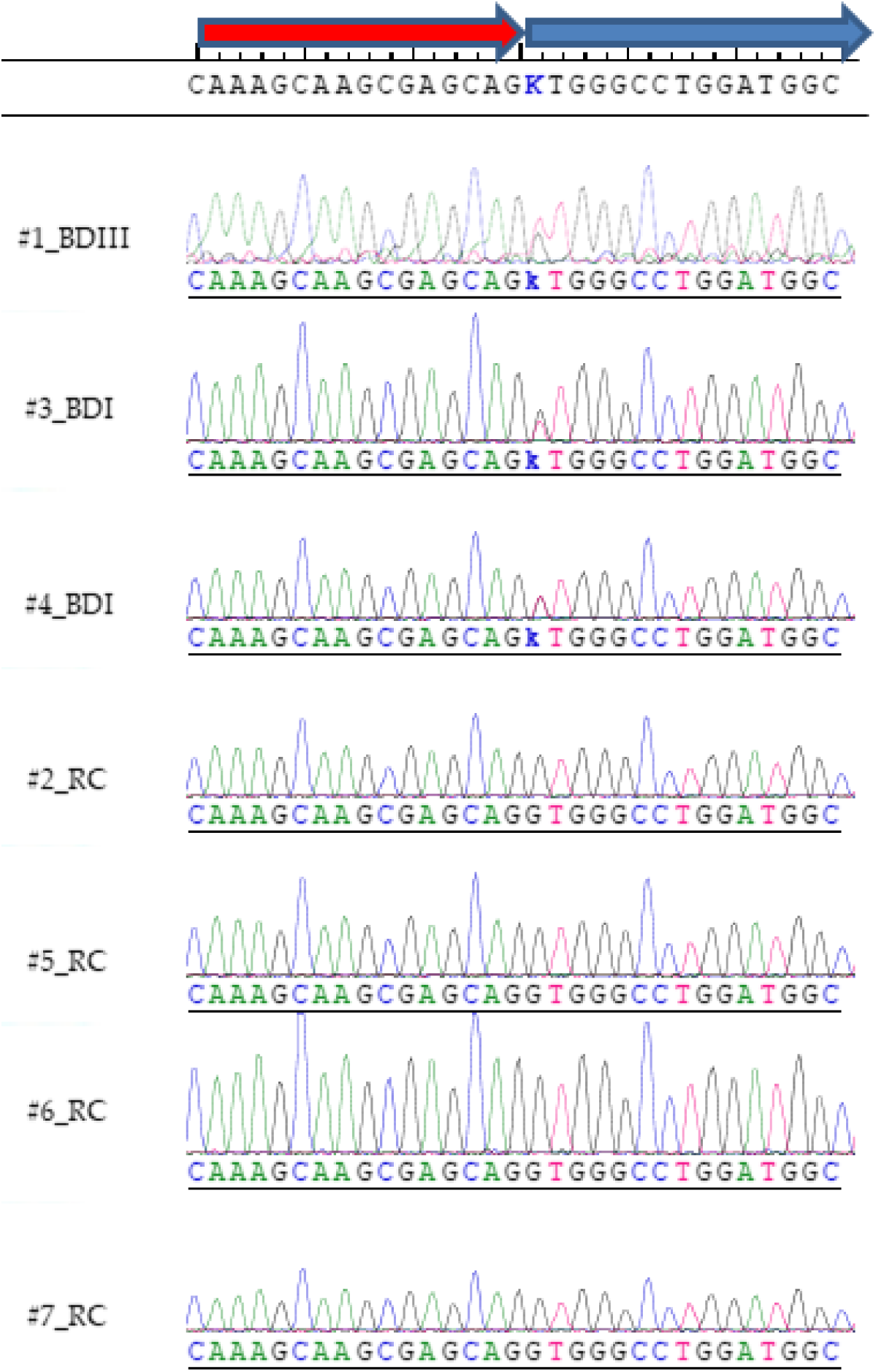
The *DCTN1* variant [NC_000002.11: g.74593585C>A (GRCh37)] was also verified using Sanger sequencing using both forward and reverse primers. Chromatograms using the forward primer are depicted. The chromatograms show heterozygous variant calls (k) for the three affected family members (Fig. 1; #1,3,4), and a normal allele for related controls (Fig. 1; #2,5,6,7). The novel variant is predicted to disrupt the invariant donor splice site (IVS22+1G>T [NM_004082.4: c.2628+1G>T]) of the *DCTN1* gene. The red arrow depicts exon 22 and the blue arrow intron 22. As the *DCTN1* gene lies on the minus strand the reverse-complement alignment is shown to highlight the GT>TT splice site variant at the invariant +1 position.

Retrograde axonal transport is the minus-end directed transport from the distal axon towards the soma and is required for neuronal autophagy. Cytoplasmic dynein is essential for retrograde axonal transport in neurons. It is the only known protein motor involved in retrograde axonal transport and transports numerous cargoes including membrane-bound organelles, mRNA, vesicles, misfolded proteins and protein aggregates. [28]. Dynactin is a large multi-protein complex which is required for dynein function including linking cargoes to dynein [29]. The *DCTN1* gene codes for the p150^Glued^ protein that is highly expressed in mammalian brain and is the largest component of the dynactin complex [30,31]. In this manuscript we specifically discuss autophagy in the context of neuronal autophagy. Rare autosomal-dominant disease-causing variants in *DCTN1* are already known for Perry syndrome (PS) [32], motor neuron disease (ALS) [33,34], and distal spinal and bulbar muscular dystrophy (dHMN7B) [35]. Psychiatric symptoms, which include severe depression, are a noted feature of PS, which is characterized by onset of atypical Parkinson disease (PD) [36]. These variants result in disruption of retrograde axonal transport and neuronal autophagy. The variants are point mutations affecting the microtubule binding region towards the N-terminus, which contrasts with our research findings in which the three affected probands are shown to possess a novel variant that is predicted to affect the C-terminus (Figs. A1, A2 in appendix). Of note, there is no historical record of PS, ALS, any muscular dystrophy, or PD in the extended family under investigation. *DCTN1* is located on 2p13.1, which is within a region previously found to show significant linkage in autosomal dominant models for BD (2p13-16) [37] and schizophrenia (2p13-14) [38]. *DCTN1* is within a region showing suggestive linkage for BD (2p11-q14) [39], as well as significant linkage for MDD (2p11.2 -p13.2) [40].

## 4. Discussion

The novel splice site variant discovered in the current research is analogous to the heterozygous dominant-negative *Gl*^*1*^ mutation in the *Drosophila Glued* gene, which is the ortholog of vertebrate *DCTN1* [41-44]. Homozygous mutations were found to be lethal in *Drosophila*. The *Drosophila* transposon-induced mutation results in a truncated transcript, which in turn results in a truncated p150^Glued^ protein missing the C-terminus (ct-p150^Glued^) [45]. The ensuing ct-p150^Glued^ fails to assemble into the dynactin complex with resulting deficits in retrograde axonal transport and neuronal autophagy [46,47]. *Glued Gl*^*1*^ mutations result in a neurodegenerative rough-eye phenotype and cause deleterious central nervous system effects in *Drosophila*, disrupting synapse formation and maturation [41]. This finding is consistent with research demonstrating that the dynactin complex is known to stabilize synapses [48]. The novel *DCTN1* splice site variant discovered in the current research is predicted to result in a PTC and a truncated protein missing the C-terminal region (Fig. A1, A2 in the appendix). Further evidence from recent structural biology work on mammalian dynein/dynactin complexes confirms that ct-p150^Glued^ would be unable to assimilate into the dynactin complex as the C-terminus normally anchors p150^Glued^ to the rest of the dynactin complex (Fig. 3) [49].

**Figure 3.**
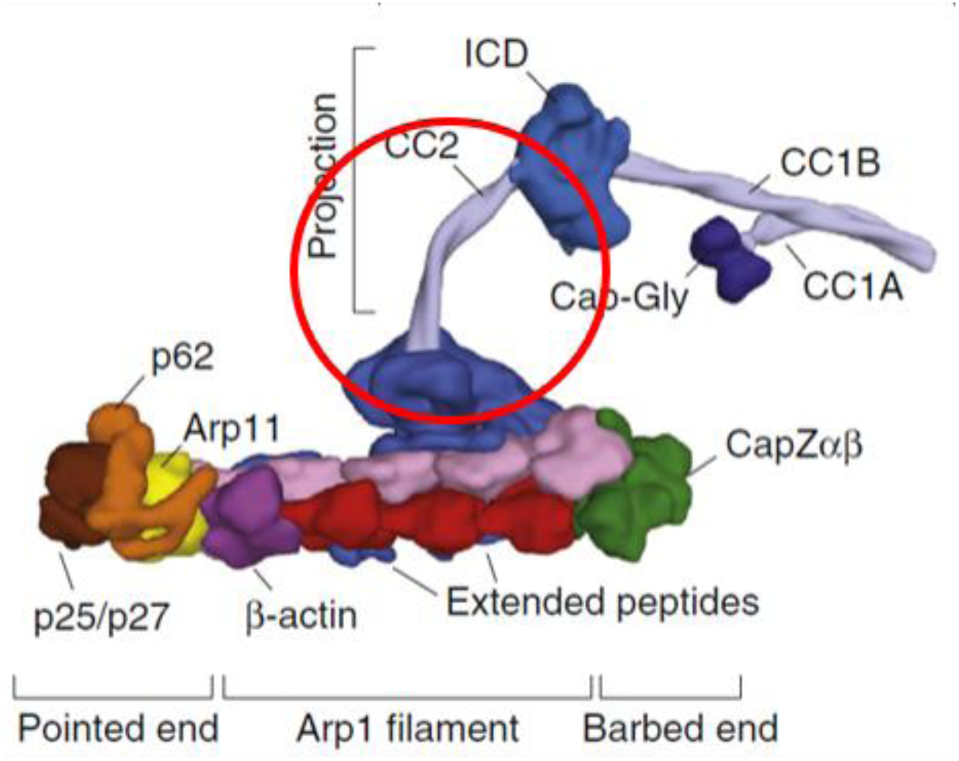
The structure of mammalian dynactin illustrating the region (circled in red) predicted to be missing as a result of the splice site variant. The long projection of p150^Glued^ is anchored to the rest of the dynactin complex by the C-terminal domain. Thus, ct-p150^Glued^ would be unable to assimilate into the dynactin complex as it lacks this domain. In contrast, disease-causing variants responsible for PS and ALS are toward the N-terminal and primarily affect the region around the Cap-Gly microtubule-binding domain. Modified from [49] with permission by the authors and Publisher (Copyright Elsevier).

If ct-p150^Glued^ is indeed produced in humans, it would be contingent on the truncated *DCTN1* transcript escaping NMD. As the PTC lies internally and is >55nt from the last exon junction it should be a very strong candidate to trigger NMD [50]. However, a ct-p150^Glued^, similar in size to the one predicted to result from the splice site variant in our current research, has previously been documented in human specimens, thus, implying that alternative splicing in this region of *DCTN1* may be under dynamic control. Fujiwara et al. discovered substantial amounts of ct-p150^Glued^ in all investigated brain specimens obtained from Alzheimer disease (AD) patients (10-70% of total p150^Glued^) [51]. ct-p150^Glued^ was not readily observable in normal brain [51]. Although their sample size was small (n = 3 AD brains), the finding of Fujiwara et al. does, nonetheless, importantly confirm that a ct-p150^Glued^ can exist in substantial amounts *in vivo*, and may represent an important feature of neurodegenerative disorders. AD is also strongly associated with psychiatric disorders with psychosis being a common event [52]. Conversely, BD patients have a significantly higher risk of developing dementia later in life [53].

As a relationship between glutamate excitoxicity and AD has been widely reported [54], Fujiwara et al suspected that this may be the cause of the observed accumulation of ct-p150^Glued^ in AD brains. They subsequently demonstrated that glutamate excitotoxicity in primary neuronal cultures similarly resulted in the production of ct-p150^Glued^, strongly suggesting a causal relationship [51]. As was the case for *Drosophila*, this truncated protein was unable to assemble into the dynactin complex and interact with cargoes (β-amyloid protein in the case of human ct-p150^Glued^), and thus represents a loss of function, disrupting retrograde axonal transport and neuronal autophagy [55]. Glutamate excitotoxicity is coupled with increased intracellular calcium concentrations [56], which are an important negative regulator of NMD [57]. Therefore, it is quite likely that formation of ct-p150^Glued^ is related to glutamate/calcium-induced impairment of NMD. Similarly, and with reference to this current research, disturbances of excitatory glutamatergic signaling are also implicated in affective disorders [58]. Indeed, many prophylactic drugs used to treat BD patients are known to modulate glutamatergic signaling [59]. BD patients in the current research cohort may be particularly sensitive to the effects of glutamatergic reinforcement of missplicing in this region of the *DCTN1* gene.

A *DCTN1* mRNA transcript of very low abundance is known (Ensembl transcript ID: DCTN1-210, ENST00000434055.5), which reflects low-level alternative splicing. This transcript is the result of alternative splicing at the identical splice junction identified in our current research and results in the same PTC. The *DCTN1* exon 22/intron 22 region has characteristics that are common to many alternatively spliced regions, namely short exons with short flanking introns [60] and G/C-rich regions typical of *cis*-regulatory elements [61] (Fig. 2). This transcript is also predicted to generate a similar sized ct-p150^Glued^ and/or trigger NMD of the truncated transcript [62]. Possibly, tissue-specific ct-p150^Glued^ is normally produced and maintained at low levels by NMD, with its formation being dependent on transient neuroexcitatory factors. Fujiwara et al. documented the presence of ct-p150^Glued^ in AD brain tissue but did not suspect alternative splicing as a possible mechanism [51,55]. However, as we have noted above this may be related to glutamate/calcium-induced impairment of NMD. Likewise, this may also occur in BD by the same mechanism and occur even without the presence of damaging splice variants in the *DCTN1* gene.

For the *DCTN1* splice site variant, and the predicted resulting formation of ct-p150^Glued^ and/or NMD, to be considered with any seriousness as being causally related to BD it would have to satisfy a major requirement: it would need to be strongly correlated with BD prophylactic drugs, antipsychotics, and antidepressants, all of which are commonly used in the treatment of BD. As discussed above, impaired retrograde axonal transport and neuronal autophagy are already known to be associated with ct-p150^Glued^. Similarly, a depletion of *DCTN1* mRNA transcripts by NMD would most likely downregulate p150^Glued^ expression with autophagic deficits [63,64]. In this regard, previous ground-breaking research has definitively linked BD prophylactic drugs to neuronal autophagy [65] The BD prophylactic drugs lithium, valproate, and carbamazepine have all been shown to enhance neuronal autophagy and thus would compensate for any deficits that are predicted to result from this variant. They achieve this by a common mechanism, namely via negatively regulating phosphoinositide signaling through depletion of inositol and by reducing *myo*-inositol-1,4,5-trisphosphate (IP_3_) [65,66]. Apart from the aforementioned psychotropic agents, the majority of antipsychotics (with two notable exceptions, namely being the typical antipsychotic haloperidol and the atypical antipsychotic clozapine) [67-69] as well as antidepressants [70,71] are also known to enhance neuronal autophagy. In addition, the autophagy-inducers rapamycin and trehalose have both been shown to have therapeutic effects in rodent models of affective disorders [72,73]. All the previously mentioned compounds are also under active consideration as therapeutic options for use in neurodegenerative disorders, by their ability to enhance retrograde axonal transport and neuronal autophagy, and to enhance clearance of aggregated and toxic protein [68,74]. Axonal transport, and associated neuronal autophagy, is known to decline with age with two distinct periods of decline, in early adulthood and later in life in rodent studies [76]. Moreover, these findings correlate with peaks for the age of onset for BD and MDD alluded to previously, inferring a relationship between decreased autophagy and the disorders.

At least two different mutant mouse models are known which suggest that deficits in neuronal autophagy may be of critical importance in the etiology of affective disorders. Reduced expression of the lysosomal aspartyl protease cathepsin D (CTSD) is known to impair neuronal autophagy [77]. In this regard, heterozygous CTSD-deficient mice exhibit BD characteristics, such as mania-related behavior and stress-induced depression [78]. Importantly, chronic administration of the BD prophylactic drugs, lithium and valproate, were found to ameliorate their symptoms and thus strongly suggest a causal relationship. In a similar manner to CTSD-deficient mice, knockdown of *DCTN5* in a murine model of psychiatric disorder susceptibility genes also results in abnormal mania-related hyperactivity [79]. *DCTN5* codes for the p25 subunit of dynactin and thus like p150^Glued^ it is intimately involved in retrograde axonal transport and neuronal autophagy [80]. It is also considered a susceptibility gene for BD [81,82]. Furthermore, deficits in neuronal autophagy are also common in lysosomal storage disorders [83], with psychiatric symptoms noted in many of these disorders [84]. For example, mania and/or psychosis has been noted in the adult forms of metachromatic leukodystrophy [85], Niemann-Pick type C disease [86] and Tay-Sachs disease [87].

Protein kinase C (PKC) has often been suggested as playing a key role in BD, as both lithium and valproate inhibit PKC signaling [88]. Thus, inhibitors of PKC have been considered as potential therapeutic agents [89]. In this regard, the selective PKC inhibitor chelerythrine, and the anti-estrogenic PKC inhibitor tamoxifen, have both been shown to have anti-manic effects in a murine model of mania [90]. Tamoxifen has also been shown to be effective in human trials for the treatment of mania [91]. PKCs are also known to suppress autophagy [92], and that includes *Drosophila* atypical PKC (aPKC) [93]. aPKCs differ from other PKC isozymes in that they are not dependent on calcium or diacylglycerol for their activity, but can nonetheless be activated by IP_3_ [94]. The neurodegenerative phenotype resulting from the *Drosophila GLUED Gl*^*1*^ mutation (which results in ct-p150^Glued^) has been shown to be rescued by mutant alleles of aPKC, thus implying that factors that negatively regulate aPKC activation (which includes reduced IP_3_ levels) alleviate the mutant phenotype [95]. Given the analogy with the observations of ct-p150^Glued^ in AD brain samples discussed previously, and its predicted formation in the BD research cohort, this finding may provide some indication of the biochemical mechanisms involved in the beneficial effects of therapeutic interventions which negatively regulate IP_3_ signaling and PKC.

The ct-p150^Glued^ predicted to occur in the current research contains a newly exposed C-terminal sequence, with a typical SH3 polyproline II helix binding motif (Pro-Pro-Pro-X-X-Pro) (Fig. A1 in appendix). Proline-rich regions such as these play an important role in protein-protein interactions which are central to cellular signaling [96]. This suggests that ct-p150^Glued^, or a cleavage product thereof, may have a physiological regulatory role, quite apart from the expected loss of function related to axonal transport. Pro-Pro-Pro-NH_2_, in a polyproline II helix conformation, has been shown to be an allosteric activator of dopamine D2 receptors, maintaining them in a high affinity state [97,98]. Thus, it is possible that ct-p150^Glued^ may represent a gain of function, activating dopamine D2 receptors in a similar manner. As dopamine D2 receptors are positive regulators of autophagy [99] this gain of function may represent a compensatory mechanism to maintain neuronal autophagy. Interestingly, a convergent functional genomics study had previously linked increased *DCTN1* gene expression with highly delusional states [75] and may be related to this proposed compensatory mechanism. The importance of dopamine in bipolar psychosis has been a central theme in the dopamine hypothesis of bipolar disorder for many decades and this proposed mechanism would be consistent with that hypothesis [100]. Dopamine D2 receptors in particular have been of intense interest as they are a common target of antipsychotic agents, and pathways leading to psychosis have been suggested to converge via high affinity state D2 dopamine receptors [101].

There is a strong likelihood that a transversion at the invariant +1 position of the splice donor site would disrupt splicing, as evidenced by previous examples of splice donor site transversions resulting in a disease phenotype (e.g. splice site disruption leading to hereditary thrombocythemia [102], in congenital afibrinogenemia [103], and mutations in CHD7 leading to CHARGE syndrome [104]). However, further research is required to validate the exact physiological effects of the splice site variant discovered in the current research cohort, and the relative contributions of NMD and ct-p150^Glued^ formation. Of note is the fact that NMD demonstrates widely differing heritable efficiencies [105], and also tissue-specific differences [106]. This may explain, at least in part, the phenotypical differences noted in the research cohort. The exon/intron 22 region of the *DCTN1* gene, as well as factors that affect splicing efficiency, and alternative splicing, in this region deserve further intense scrutiny in psychiatric disorders. It would also be highly desirable to identify this splice variant in other affected families to further validate its significance and to obtain an estimate of relative frequency.

## 5. Conclusions

The *DCTN1* splice site variant discovered in the current research is not only co-segregating, novel, and in a highly evolutionary conserved region, but it is also predicted to be deleterious. Taken together all these qualities strongly suggest pathogenicity of the splice site variant. Furthermore, the neuronal autophagy deficits that are predicted to result are consistent with established knowledge regarding the therapeutic interventions used in BD and MDD. Previously discovered disease-causing variants in *DCTN1* (e.g. in PS, ALS) are inherited in an autosomal-dominant fashion, and disease-causing variants in PS (resulting in atypical PD) are already known to also result in severe depression. Thus, there is a strong likelihood that the variant uncovered in the present research may also reflect a primary autosomal dominant cause of psychiatric pathology in the research cohort.

As noted previously, a ct-p150^Glued^, which is predicted to result from the splice site variant discovered in this research, has also been found in AD brain samples as well as in *Drosophila* mutants. In both cases it is related to deficits in neuronal autophagy. Furthermore, deficits in neuronal autophagy are implied as being causal in at least two different murine models of affective disorders. As also noted previously, axonal transport and associated neuronal autophagy similarly slow at ages that are likewise consistent with onset of illness in both BD as well as recurrent MDD. In addition, lysosomal storage diseases, which are characterized by deficits in neuronal autophagy, are also associated with psychiatric illness. Therefore, this novel co-segregating splice site variant, which is predicted to impair neuronal autophagy, may also hold an important key to the understanding of the pathological biochemical mechanisms involved in BD and MDD in general.

In summary, the proposed mechanisms are highly speculative, but nonetheless they form a framework for future research in many areas of neuropsychiatric and neurodegenerative diseases.

## Abbreviations

AD: Alzheimer disease
ALS: motor neuron disease
aPKC: atypical PKC
BDI: bipolar disorder I
BDII: bipolar disorder II
BDIII: bipolar disorder III or cyclothymic disorder
ct-p150Glued: C-terminal truncated p150Glued protein
*DCTN1*: dynactin-1 gene
IP3: *myo*-inositol-1,4,5-trisphosphate
MDD: major depressive diagnosis
NMD: RNA nonsense-mediated decay
NGS: next generation sequencing
PD: Parkinson disease
PKC: protein kinase C
PS: Perry syndrome
PTC: premature termination codon
WES: whole exome sequencing.

## Supplementary Materials

S1, variant calls; S2, Sanger reads. The following are available online at www.mdpi.com/xxx/

## Author Contributions

AH initiated this research project, was responsible for experimental design, and performed all analyses. AH and AJLC wrote and edited the manuscript. All authors have read and agreed to the published version of the manuscript.

## Acknowledgments

Prof. Ole A. Andreassen (Psychosis Research Centre, Oslo University, Norway) and Dr. Srdjan Djurovic (Dept. Medical genetics, Oslo University Hospital, Norway) are thanked for sample collection from unaffected controls, subsequent genomic DNA extraction and assembly of informed consent forms, as well as shipment of the relevant DNA samples for subsequent sequencing. Andrew Carter (U. Cambridge, UK) is thanked for permission to use his graphic depicting the structure of the dynactin complex. The following are thanked for providing valuable feedback and/or proof-reading during the preparation of the manuscript: Jenny Vo (Teva Pharmaceuticals, Australia), Jan Fullerton (UNSW, Australia), Steve Ambrose (Ambrose Ecological, Australia), Philip Kuchel (U. Sydney, Australia), Fei Liu (Macquarie U., Australia), Daria Mochly-Rosen (Stanford U., USA), and Andrew Carter (U. Cambridge, UK). Guy Plunkett III and Eric Ma are thanked for providing DNASTAR Lasergene software technical support (DNASTAR, Madison, Wisconsin, USA).

## Appendix A

**Figure A1.**
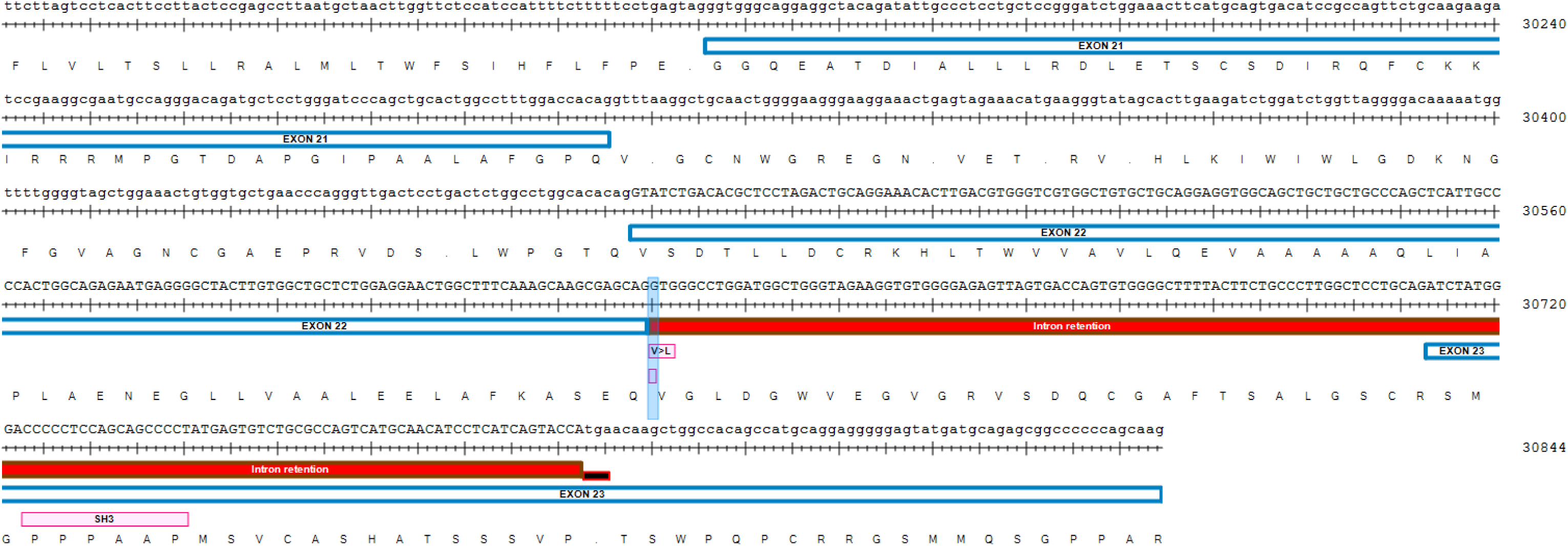
(following page) The region of the *DCTN1* gene illustrating the splice site variant (NG_008735.2: g.30630G>T) which is predicted to disrupt the invariant +1 splice donor site, resulting in intron retention and a premature termination codon (PTC). The transcript can be degraded by NMD and/or be translated to form a C-truncated p150^Glued^ protein. The exons/introns in this region are short which is characteristic of alternatively spliced regions. The new C-terminus tail includes an amino acid variant with a predicted benign change V>L (V877L) and is characterized by a SH3 polyproline II helix binding motif [PPPxxP] which implies that the C-terminal truncated p150^Glued^ protein may have a physiological role which is hypothesised to involve allosteric activation of dopamine D2 receptors.

**Figure A2.**
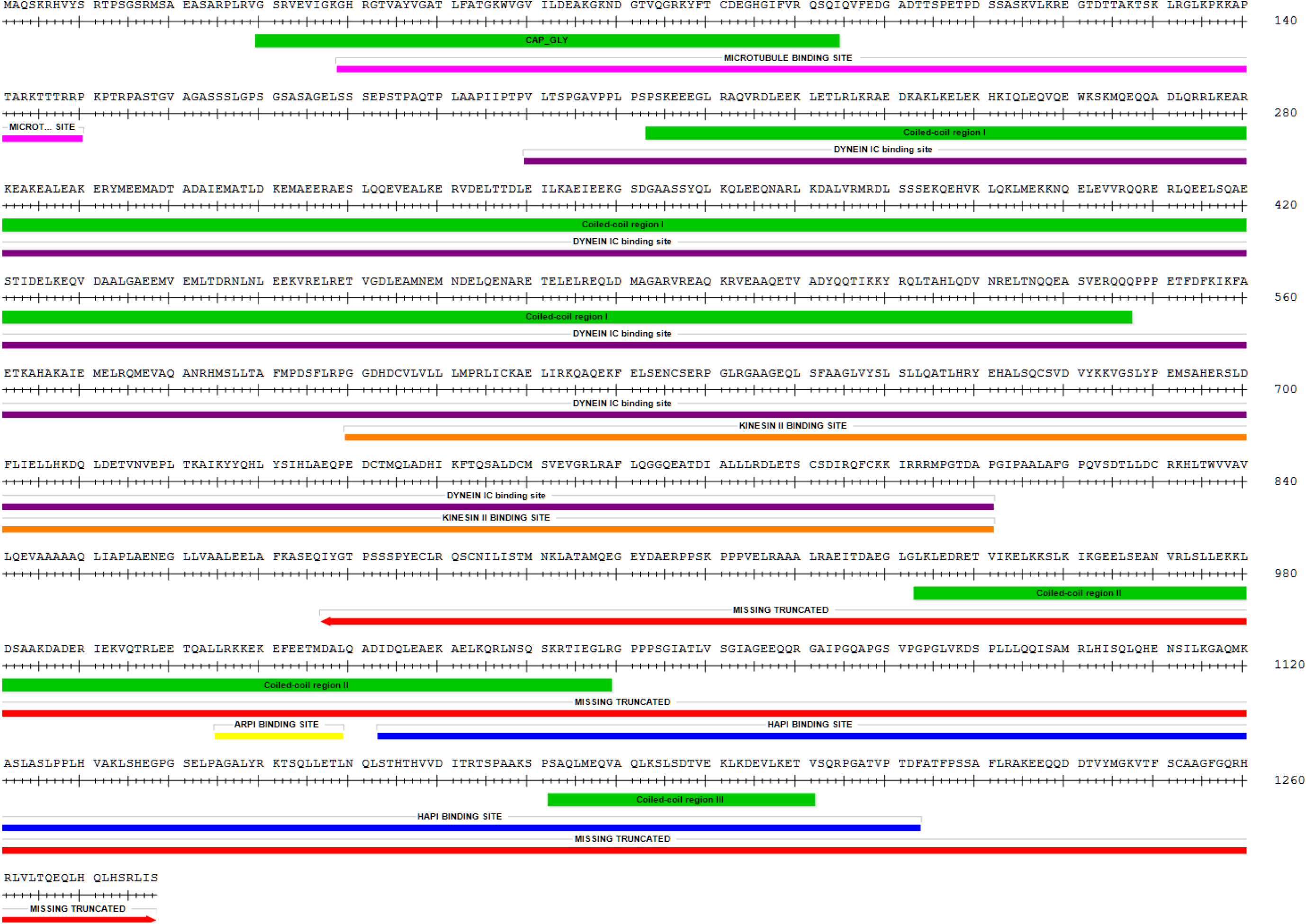
(following page) p150^Glued^ protein (UniProt ID: Q14203, 1278AA). The colored bars under the amino acid sequence illustrate the different regions: ***Red***, the region predicted in this study to be missing in ct-p150^Glued^ due to a splice site variant; ***Green***, CAP_GLY and coiled-coil domains; ***Magenta***, microtubule binding domain; ***Purple***, dynein IC binding region; ***Orange***, kinesin II binding region; ***Yellow***, ARPI binding region; ***Blue***, HAP1 binding region.

