## Supplementary material for "A novel co-segregating *DCTN1* splice site variant in a family with Bipolar Disorder may hold the key to understanding the etiology": Sanger alignment.pdf

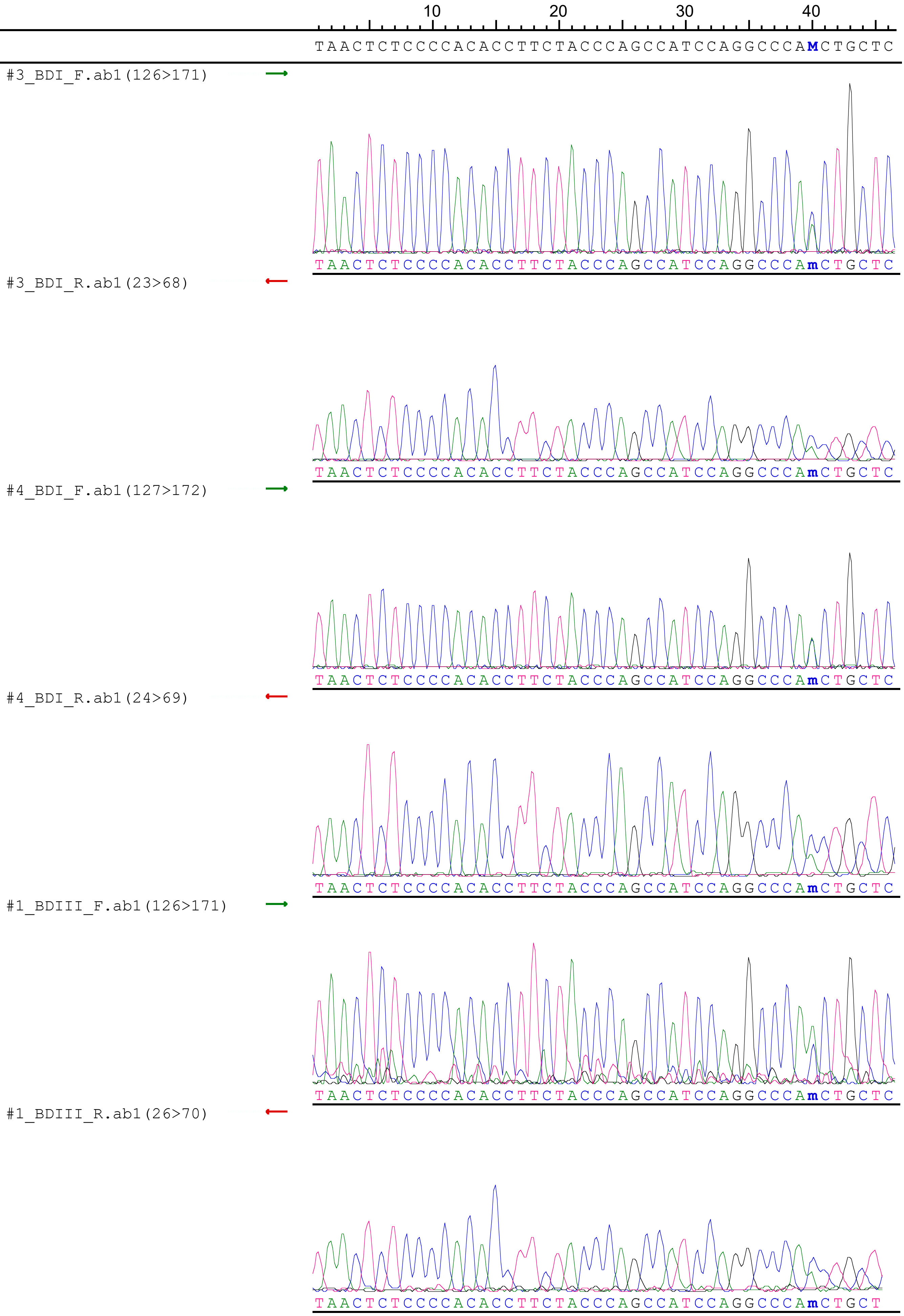
10
20
30
40
TAACTCTCCCCACACCTTCTACCCAGCCATCCAGGCCCAMCTGCTC
#3_BDI_F.ab1(126>171)
TAACTCTCCCCACACCTTCTACCCAGCCATCCAGGCCCAmCTGCTC
#3_BDI_R.ab1(23>68)
TAACTCTCCCCACACCTTCTACCCAGCCATCCAGGCCCAmCTGCTC
#4_BDI_F.ab1(127>172)
TAACTCTCCCCACACCTTCTACCCAGCCATCCAGGCCCAmCTGCTC
#4_BDI_R.ab1(24>69)
TAACTCTCCCCACACCTTCTACCCAGCCATCCAGGCCCAmCTGCTC
#1_BDIII_F.ab1(126>171)
TAACTCTCCCCACACCTTCTACCCAGCCATCCAGGCCCAmCTGCTC
#1_BDIII_R.ab1(26>70)
TAACTCTCCCCACACCTTCTACCCAGCCATCCAGGCCCAmCTGCT

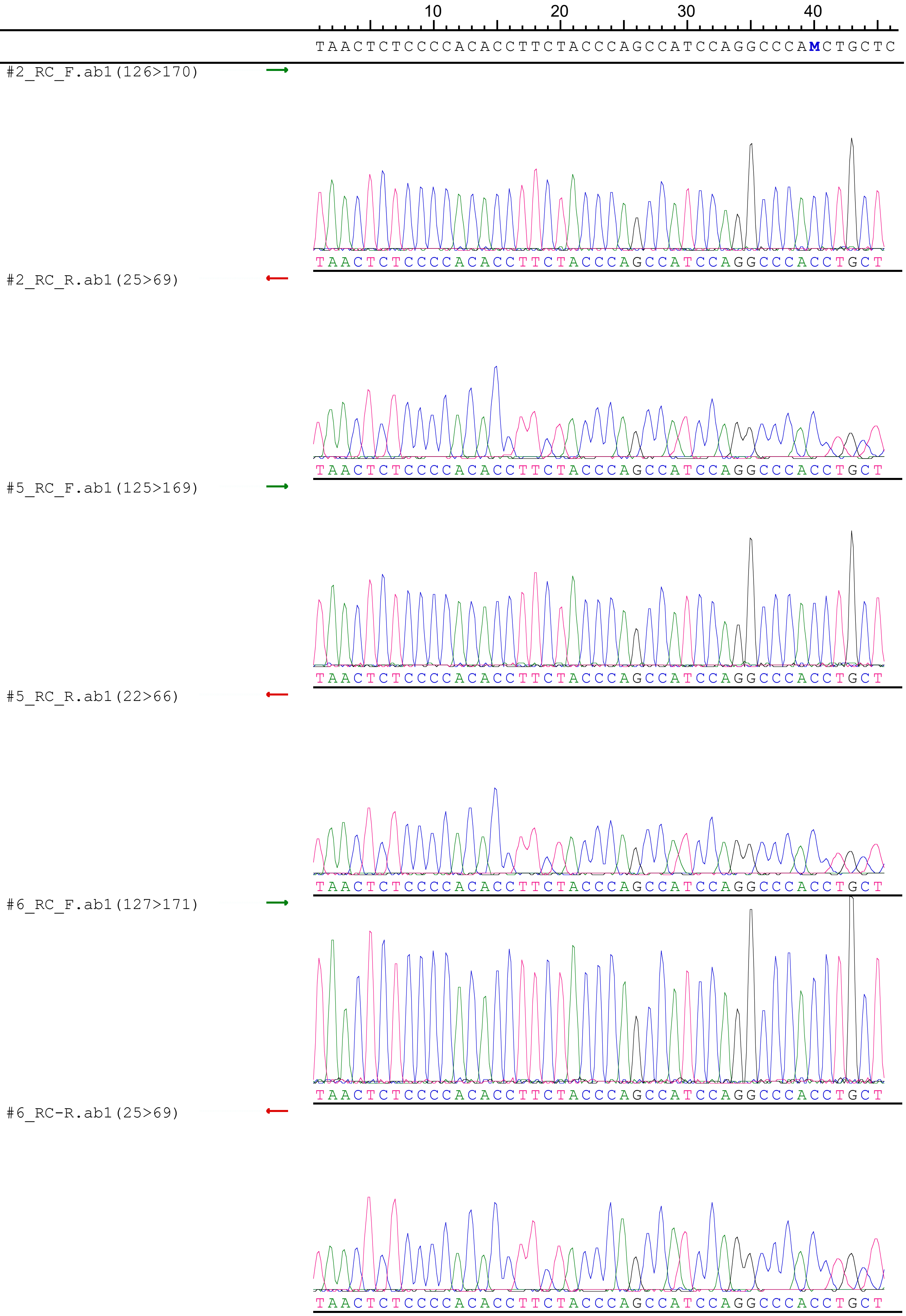

10
20
30
40
TAACTCTCCCCACACCTTCTACCCAGCCATCCAGGCCCAMCTGCTC
#2_RC_F.ab1(126>170)
TAACTCTCCCCACACCTTCTACCCAGCCATCCAGGCCCACCTGCT
#2_RC_R.ab1(25>69)
TAACTCTCCCCACACCTTCTACCCAGCCATCCAGGCCCACCTGCT
#5_RC_F.ab1(125>169)
TAACTCTCCCCACACCTTCTACCCAGCCATCCAGGCCCACCTGCT
#5_RC_R.ab1(22>66)
TAACTCTCCCCACACCTTCTACCCAGCCATCCAGGCCCACCTGCT
#6_RC_F.ab1(127>171)
TAACTCTCCCCACACCTTCTACCCAGCCATCCAGGCCCACCTGCT
#6_RC-R.ab1(25>69)
TAACTCTCCCCACACCTTCTACCCAGCCATCCAGGCCCACCTGCT

10 20 30 40

TAACTCTCCCCACACCTTCTACCCAGCCATCCAGGCCCA**M**CTGCTC

#7_RC_F.ab1(128>172)

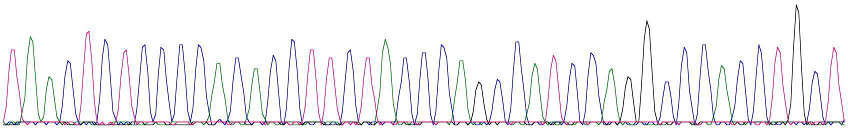

TAACTCTCCCCACACCTTCTACCCAGCCATCCAGGCCCACCTGCT

#7_RC_R.ab1(29>73)

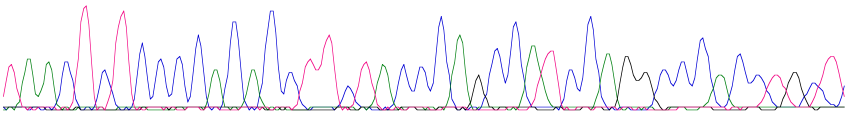

TAACTCTCCCCACACCTTCTACCCAGCCATCCAGGCCCACCTGCT
